# Deletion of core septin gene *aspB* in *Aspergillus fumigatus* results in fungicidal activity of caspofungin

**DOI:** 10.1101/2025.02.25.640155

**Authors:** Rebecca Jean Busch, Carson Doty, C. Allie Mills, Flutur Latifi, Laura E. Herring, Vjollca Konjufca, Francisco Carvallo, Tatiana Boluarte, José M Vargas-Muñiz

**Affiliations:** Department of Biological Sciences, Virginia Tech, Blacksburg, Virginia, United States; Center for Emerging, Zoonotic, and Arthropod-borne Pathogens, Virginia Tech, Blacksburg, Virginia, United States; School of Biological Sciences, Southern Illinois University-Carbondale, Carbondale, Illinois, United States; Michael Hooker Metabolomics and Proteomics Core Facility, Department of Pharmacology, University of North Carolina at Chapel Hill, Chapel Hill, North Carolina, United States; Microbiology Program, Southern Illinois University-Carbondale, Carbondale, Illinois, United States; Department of Biomedical Sciences and Pathobiology, Virginia-Maryland College of Veterinary Medicine, Blacksburg, Virginia, United States; Fralin Life Science Institute, Virginia Tech, Blacksburg, Virginia, United States

## Abstract

Septins are a family of GTP-binding proteins found in many eukaryotic lineages. Although highly conserved throughout many eukaryotes, their functions vary across species. In *Aspergillus fumigatus*, the etiological agent of invasive aspergillosis, septins participate in a variety of processes, including conidiation, septation, and responses to cell wall stress. Previous studies determined that the Δ*aspB* strain had a greater sensitivity to anti-cell wall drugs, especially the echinocandins, yet mechanisms behind this augmented sensitivity are unknown. We performed cell viability staining of the deletion strains after caspofungin exposure and found that the Δ*aspA*, Δ*aspB*, and Δ*aspC* strains had significantly lower cell viability. Concomitant with the reduced viability, deletion strains are more susceptible to caspofungin on solid media. These results indicate that the septin cytoskeleton is important for *A. fumigatus* survival in the presence of caspofungin. Due to the potential of improved therapeutic outcome, we followed up using a neutropenic murine model of invasive aspergillosis. Animals infected with the Δ*aspB* strain and treated with caspofungin showed improved survival compared to the animals infected with *akuB*^KU80^ wild-type or complemented strains. Additionally, histological analysis showed reduced fungal burden and inflammation in the Δ*aspB* infected, caspofungin-treated group.

Affinity purification coupled with quantitative proteomics identified proteins involved in the septin-dependent response to caspofungin, including four candidate interactors involved in cell wall stress response. Deletion of these candidate genes resulted in increased susceptibility to caspofungin and moderately reduced viability post-drug exposure. Taken together, these data suggest that septin AspB contributes to the fungistatic response to caspofungin.

**Author Summary:** Invasive aspergillosis is a pulmonary disease caused by the fungus *Aspergillus fumigatus* that primarily occurs in immunocompromised patients. Invasive aspergillosis has a high mortality rate, ranging from 50-90%. Therapy options are limited due to few available drugs with fungicidal activity and growing global drug resistance. Treatment typically starts with triazoles, which target the fungal cell membrane. If unsuccessful, an echinocandin, which targets the cell wall, is given as a salvage therapy in the U.S. Echinocandins, including caspofungin, are fungistatic against *A. fumigatus*, slowing growth of the fungus rather than killing it. Due to this, echinocandins have a high therapeutic failure rate. Previous work suggests that deletion of the cytoskeletal septin genes increases sensitivity to caspofungin. Here we describe our finding that the septin genes *aspA, aspB*, and *aspC* are involved in the fungal response to caspofungin.

Additionally, the deletion of *aspB* results in fungicidal activity of this otherwise fungistatic drug. These findings show promise for novel therapy options that block the septin-mediated response to caspofungin.

## Introduction

*Aspergillus fumigatus* is a ubiquitous environmental mold responsible for a wide range of opportunistic systemic and allergic pathologies [1]. One such pathology is invasive aspergillosis (IA), an invasive infection most common in immunocompromised patients [2]. Over 300,000 cases of IA are reported yearly, leading to a mortality rate ranging between 30-90% [1,3].

Additionally, the at-risk patient population for IA is increasing due to increasing numbers of immunomodulating therapies, as well as emerging global diseases [4–6]. IA has the highest per-patient cost of any invasive fungal disease, costing the United States an estimated $1.3 billion per year [7].

IA begins when the conidia, or asexual spores, are inhaled by an individual [8,9]. Due to their small size (∼2.5 μm diameter) and hydrophobic nature, the conidia can travel down the airway into the terminal alveoli [10]. In healthy individuals, the mucociliary escalator removes conidia [8]. However, any remaining conidia can then germinate within the lung. Epithelial cells, alveolar macrophages, neutrophils, and other immune cells can detect the germinating conidia, which leads to the release of cytokines and phagocytosis of conidia by immune cells [8]. However, immunocompromised patients lack the appropriate immune responses, which leads to the progression of IA. Germinated conidia can then establish polarity, leading to the formation of hyphae that invades host tissue. IA can disseminate within the host via the release of hyphal fragments into the bloodstream. Neutropenic patients, such as those on anti-rejection medications post-transplant, are at high risk for IA [11]. In neutropenic patients, disease often presents with abundant hyphal growths, angioinvasion, and intra-alveolar hemorrhage [12,13]. Patients undergoing corticosteroid-induced immunosuppression are also at risk of IA, as glucocorticoids lead to a reduction of pattern recognition receptor (PRR) signaling and inhibit lymphocyte activation [14,15]. Pneumonia, inflammatory necrosis, and minimal hyphal growths are seen in cases of IA in patients under corticosteroid-induced immunosuppression [12,13].

Therapeutic options to effectively treat IA are limited. The frontline treatment for IA is the triazole class of drugs [16]. Triazole treatments are often prolonged and are now common to use as antifungal prophylaxis in at-risk populations. Nonetheless, cases of triazole-resistant *A. fumigatus* infections are increasing in some countries and are a major concern worldwide. This increase in the incidence of triazole resistance led to the inclusion of *A. fumigatus* as a critical priority in the World Health Organization 2022 Fungal Priority Pathogens List [17].

Echinocandins are a class of antifungals that inhibit the β-glucan synthase [18]. In fungi such as *C. albicans*, echinocandins have a fungicidal effect [19]. In contrast, the echinocandins are fungistatic against *A. fumigatus* [18]. For immunocompromised patients in the United States, the echinocandins are often used as a salvage or secondary therapy in conjunction with triazoles as they cannot reap the benefits of solely fungistatic medications [16]. Slowing the growth of the fungi through fungistatic therapies then requires either the immune system or fungicidal drugs to kill the fungi and clear disease [20,21]. Because of its fungistatic nature, caspofungin use as a salvage treatment had a favorable response rate of only 45% [22]. Understanding the mechanisms within fungi that render these medications fungistatic rather than fungicidal can aid in developing new therapies that improve the efficacy of existing antifungal drugs.

Previous studies have shown that septins AspA, AspB, and AspC contribute to the *Aspergillus spp.* response to the echinocandin caspofungin [23,24]. Septins are a family of highly-conserved eukaryotic GTP-binding proteins. In *A. fumigatus,* septins are primarily involved in septation, conidiation, and response to cell wall stress [24]. *A. fumigatus* has five septins: AspA, AspB, AspC, AspD, and AspE [25]. AspA-D are core mitotic septins involved in polymerizing into hexameric and octameric complexes. They are orthologs of *Saccharomyces cerevisiae*’s Cdc11, Cdc3, Cdc12, and Cdc10, respectively [26]. AspE is a Momany Group 5 septin that is present in filamentous fungi and other eukaryotic lineages [27]. *Aspergillus nidulans* septins have been associated with the cell wall integrity (CWI) pathway [23]. Double deletion strains of *aspB* and CWI pathway MAPK *mpkA* showed a novel phenotype when grown on caspofungin compared to the respective single deletion strains [23]. In *Candida albicans,* deletion of septin genes leads to aberrant localization of chitin in the cell wall, increasing susceptibility to caspofungin [28,29]. This susceptibility is due to caspofungin inducing aberrant cytokinesis and phosphatidylinositol (4,5)-bisphosphate relocalization [28–30]. This points out a conserved role of septins in the fungal response to caspofungin, but the mechanism might not be conserved.

In this study, we found that septins AspA, AspB, and AspC contribute to fungal viability post-caspofungin exposure in *A. fumigatus*. The deletion of *aspB* in particular produced a fungicidal response. We also observed a higher chance of survival, reduced lung inflammation, and reduced fungal burden were associated with Δ*aspB* infected mice treated with caspofungin than Δ*aspB* strain treated with saline or *akuB*^KU80^ and Δ*aspB::aspB* strains with either treatment in our neutropenic murine model of IA. To gain deeper mechanistic insights, we conducted a quantitative mass spectrometry-based proteomics analysis to identify candidate AspB-interacting proteins during exposure to caspofungin. Based on our proteomics analysis, we narrowed down six possible proteins involved in the fungal response to caspofungin that were significantly increased (by at least two-fold) upon caspofungin exposure. Gene ontology (GO) analysis indicated that these candidate protein interactors might have a role in cell wall function and organization, suggesting that AspB may mediate cell wall responses through its interactions with these candidates. Deletion strains for each candidate were generated and characterized to obtain an understanding of their biology at basal conditions. Strains were then tested against caspofungin, and four out of the six genes, *bgt1, gel2, nsdD*, and *mapA*, were implicated in fungal viability after caspofungin exposure. Double deletion mutants lacking *aspB* and either *bgt1*, *gel2*, *nsdD*, and *mapA* were generated, which indicated that their role in caspofungin response is mediated through AspB.

## Results

### Deletion of *aspA*, *aspB*, and aspC Reduces Viability Post-Caspofungin Exposure

Previous work demonstrated that the *A. fumigatus* Δ*aspB* and Δ*aspC* strain had increased sensitivity to caspofungin on solid media using single-spot inoculation [24]. Although informative, these assays provide a minimal understanding of the effect of caspofungin in the septin deletion strains. In order to further characterize the role septins in *A. fumigatus* response to echinocandins, we conducted a spore dilution assay (10^4^-10^1^ conidia, 1 µg/mL caspofungin), minimum effective concentration assay (MEC), and E-strip test (10^6^ conidia) to visualize the susceptibility to caspofungin. Deletion of the core septin genes, *aspA, aspB*, and *aspC*, resulted in increased sensitivity to caspofungin compared to the *akuB*^KU80^ wild-type strain, Δ*aspB::aspB* complemented strain, and other septin deletion strains in these assays (Fig. 1A, Supplementary Table 4). Similarly, the Δ*aspA*, Δ*aspB*, and Δ*aspC* strains had a clearer zone of effect on the E-strip test (Fig. 1D). To explore whether this effect was exclusive to caspofungin, we conducted an E-strip test to visualize the susceptibility to another drug in the echinocandin class, micafungin. The Δ*aspA and* Δ*aspC* strains showed a slight increase in susceptibility to micafungin, as seen by a clearer zone of effect, while Δ*aspB* had the most noticeable decrease in growth in the zone of effect (Fig. 1D). Thus, this increase in susceptibility of the Δ*aspA,* Δ*aspB*, and Δ*aspC* strains is not limited to caspofungin, albeit it is more pronounced.

**Figure 1.**
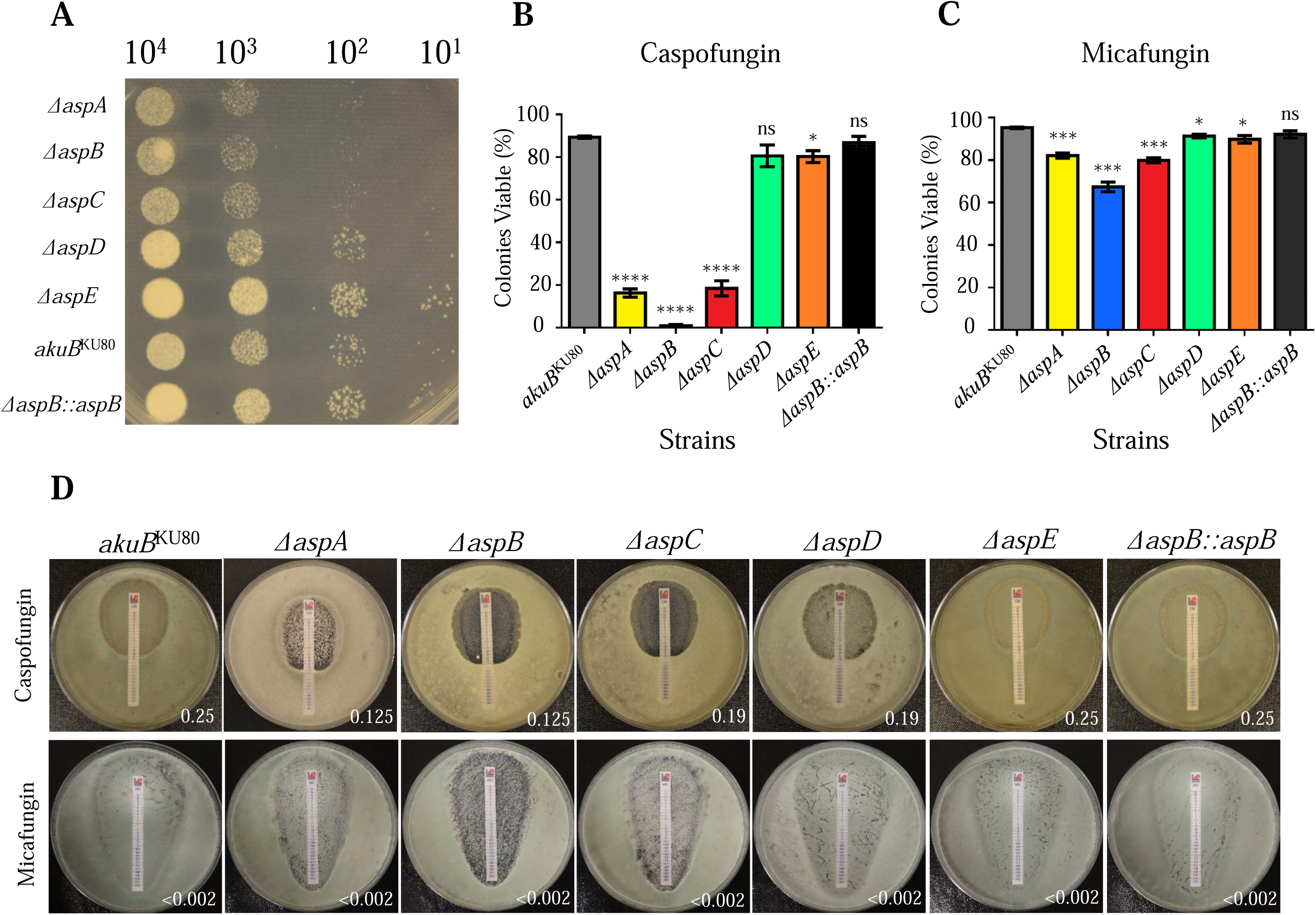
AspA, AspB, and AspC are involved in fungal response to caspofungin. The deletion of *aspA, aspB, and aspC* results in reduction of viability to echinocandins. (A) Spore dilution assay show an increase in susceptibility to caspofungin in the Δ*aspA,* Δ*aspB,* and Δ*aspC* strains. 10^4^-10^1^ conidia were plated on GMM media supplemented with caspofungin for 48 hours at 37°C. (B, C) Deletion of *aspB* results in loss of viability when grown in caspofungin but not micafungin. 10^4^ conidia were incubated for 48 hours in GMM supplemented with (B) 1µg/mL caspofungin or (C) 1 µg/mL micafungin. Cells were incubated in CFDA for one hour then visualized. Viable and non-viable colonies were counted. The number of viable colonies was divided by the total number of colonies. Experiment was replicated three times. Error bar represent SEM. Student t-test were done in Graphpad Prism, with each strain compared against the *akuB*^KU80^ wild-type. Asterisks denote statistical significance levels as follows: ns p>0.05, * p≤0.05, ** p≤0.01, *** p≤0.001, and **** p≤0.0001. (D) E-strip plates of caspofungin and micafungin show clearer zone of effect in Δ*aspA,* Δ*aspB*, and Δ*aspC* strains. 10^6^ conidia were plated with beads and left to dry. E-strip was placed, and plates were grown at 37°C for 48 hours. All experiments were replicated three times. Representative images are shown in this figure.

### AspB is Required for Fungistatic Response to Caspofungin *in vitro*

Since we observed an increase in susceptibility in all three of our drug susceptibility assays, we hypothesized that the absence of AspA, AspB, and AspC leads to a fungicidal effect of caspofungin (Fig. 1). To determine the viability of the deletion strains during caspofungin exposure, we utilized CFDA (5,(6)-Carboxyfluorescein Diacetate). When CFDA is cleaved by esterases, its product is fluorescent, allowing us to use it as a way to distinguish between viable and non-viable cells [31]. In basal conditions, 10^4^ conidia were grown on coverslips in GMM+UU (glucose minimal media + uracil and uridine) liquid media for 24 hours, followed by 1 hour incubation in 1 µg/mL CFDA. Uracil and uridine are added to the media due to Δ*aspD* being auxotrophic. All strains had equally viable confluent growth, indicating that the deletion of the septin does not impact mycelial viability(Fig. 2A). Strains were then grown for 48 hours in the presence of 1 µg/mL caspofungin, followed by incubation with CFDA. We then quantified the number of viable and total microcolonies formed in the presence of caspofungin. The *aspA, aspB*, and *aspC* deletion strains were significantly less viable (p<0.0001) after 48 hours of caspofungin exposure, with only 0.92% of Δ*aspB* (p<0.0001) being viable (Fig. 1B, 2B). Δ*aspB* also does not form many microcolonies in the presence of caspofungin. Δ*aspA* and Δ*aspC* form microcolonies in the presence of caspofungin; however, only 16.3% and 18.5% of the colonies, respectively, were viable at the time of microscopic examination (p<0.0001)(Fig. 1B, 2B). We repeated this assay with the echinocandin micafungin in order to test if this reduction in viability was specific to caspofungin or a general response to echinocandin exposure. In contrast to caspofungin treatment, the majority of colonies from all strains were viable after micafungin treatment (*akuB*^KU80^=95.3%, Δ*aspA*=82.1%, Δ*aspB*=67.4%, Δ*aspC*=79.9%, Δ*aspD*=91.4%, Δ*aspE*=89.8%, Δ*aspB::aspB*=92.2% viable colonies)(Fig. 1C). This indicates that the fungicidal effect in the Δ*aspB* strain is specific to caspofungin.

**Figure 2.**
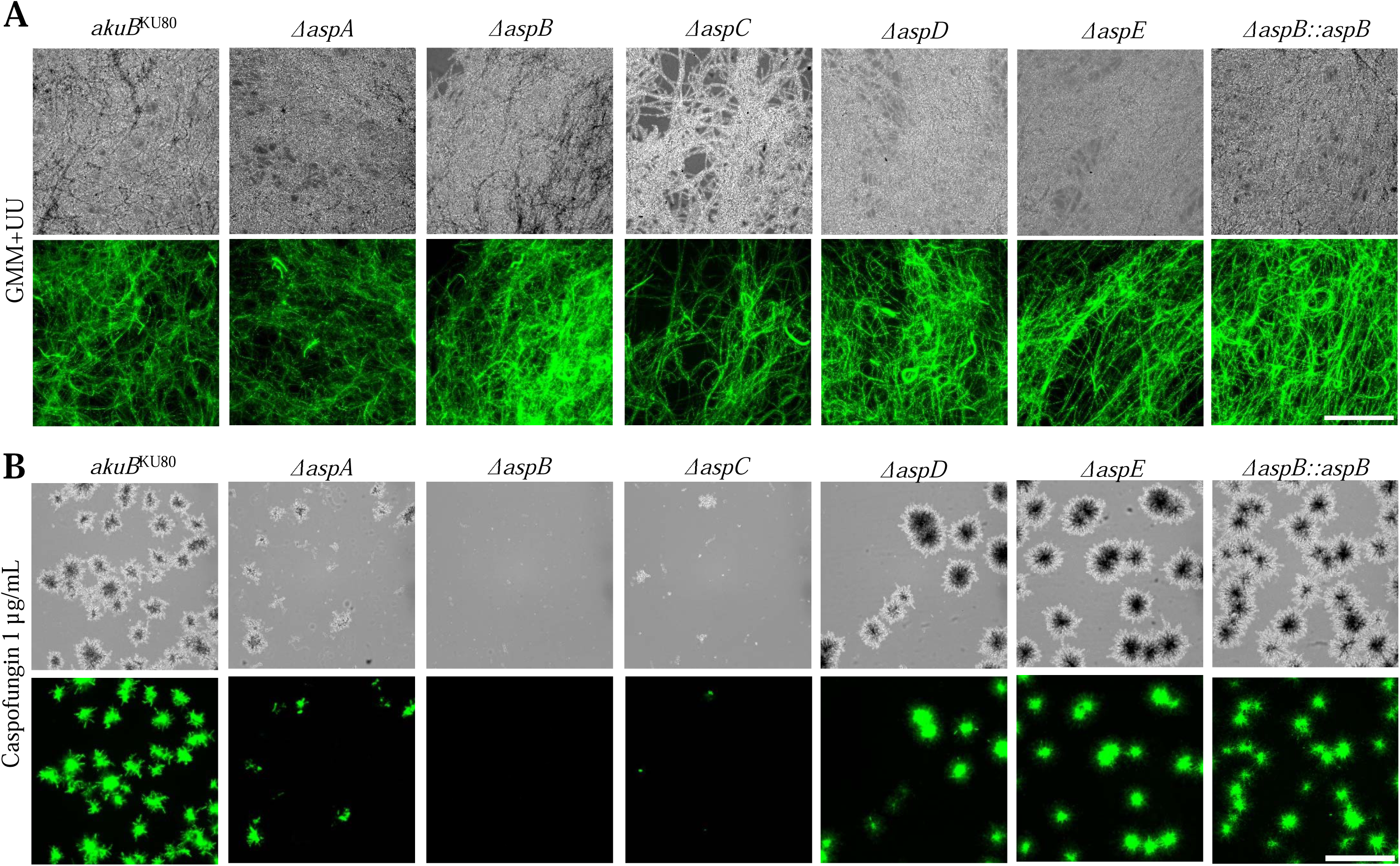
Δ***aspB* strain loses cell viability when grown in caspofungin.** (A) All strains had viability when grown in GMM+UU. 10^4^ conidia were grown in 4 mL GMM+UU at 37°C for 24 hours. Cells were then treated with CFDA, which is hydrolyzed in living cells to a fluorescent ester, then visualized. (B) Loss of viability in Δ*aspB* strain is seen after growth in the presence of caspofungin. After 48 hours post exposure to 1 µg/mL caspofungin, cells were incubated in CFDA for one hour then visualized. Experiments were replicated three times. Scale bar is 500 µm.

### AspB is Involved in Response to Caspofungin in Mature Mycelium

It is possible that most of these phenotypes that we observe are due to caspofungin acting as the spore germinates, as exemplified by the small microcolonies in the Δ*aspB* strain. Mature mycelia would be a more clinically relevant growth stage, and for this reason, we decided to determine the effect of caspofungin on mature mycelia. To test this, we used propidium iodide (PI) stain to determine hyphal damage. Strains were grown in GMM+UU for 24 hours at 37°C before being stained with PI and visualized. No hyphal damage was seen under basal conditions (Fig. 3A). Following the same procedure, we grew strains then incubated them with caspofungin (1 µg/ml) for 2 hours. Coverslips were washed and stained as before, then visualized. Similar to our viability assays, only the Δ*aspB* strain exhibits extensive hyphal damage after 2 hours of exposure to caspofungin (Fig. 3B). Taken together, these results suggest that AspB is needed for fungal response to caspofungin.

**Figure 3.**
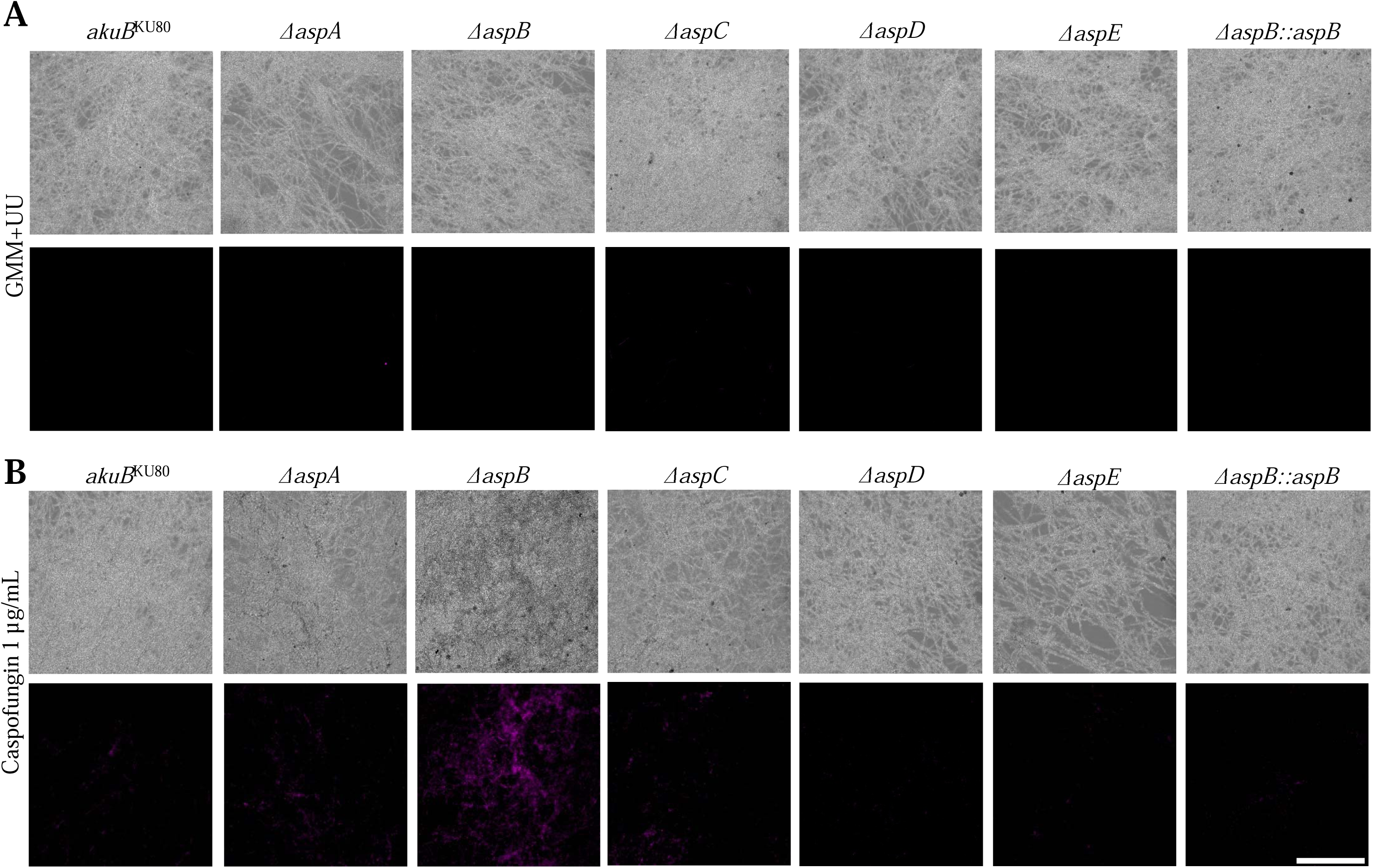
Δ***aspB* strain has increased hyphal damage during caspofungin exposure.** (A) No hyphal damage seen in basal conditions. 10^4^ conidia were grown in GMM at 37°C for 24 hours. Cells were then washed with PIPES (pH 6.7) for 5 minutes twice. Slides were then treated with propidium iodide (PI) solution, which stains nucleic acids. Coverslips were washed twice with PIPES, then prepared and visualized. (B) Hyphal damage was seen in Δ*aspB* strain treated with caspofungin. 10^4^ conidia were grown in GMM at 37°C for 24 hours. Coverslips were then incubated in caspofungin for 2 hours at 37°C. Cells were then washed with PIPES and treated with PI solution. Experiments were replicated three times. Scale bar is 500 µm.

### Deletion of *aspB* improves caspofungin treatment efficiency in a neutropenic murine model of invasive aspergillosis

As we observed a strong fungicidal effect against the Δ*aspB,* we decided to determine if deletion of *aspB* led to improved survival in our murine model of invasive aspergillosis. As the Δ*aspB* strain has been previously shown to have no discernible difference in virulence compared to the *akuB*^KU80^ and Δ*aspB::aspB* strains with respect to fungal burden and lung inflammation, any effect on survival would be attributed to the role that AspB plays in fungal response to caspofungin [24]. To test this hypothesis, neutropenia was induced in 6-week old male CD-1 mice using 175 mg/kg cyclophosphamide and 40 mg/kg triamcinolone acetonide. Neutropenic mice were then intranasally infected with 4×10^6^ conidia of *akuB*^KU80^, Δ*aspB*, and Δ*aspB::aspB* strains. They were subsequently treated with either 2 mg/kg caspofungin or an equivalent volume of the saline vehicle. All groups of mice treated with saline all had the first death on day 3 post-infection (Fig. 4A) and were not statistically significantly different. Mice inoculated with Δ*aspB* strain and treated with caspofungin had a 70% survival rate (p<0.0001)(Fig. 4B). Additionally, in this group the first death did not occur until day 12 post-infection. Compared with the next highest groups, the Δ*aspB* strain treated with saline and *akuB*^KU80^ treated with caspofungin had 30% survival (Fig. 4A). Lung histology was then performed to visualize inflammation and fungal lesions. Concurrent with the survival graph, animals infected with the Δ*aspB* strain treated with caspofungin had stunted fungal growth compared to animals from the other groups, while inflammation around fungal cells was similar across conditions (Fig. 4D). Previous infection models also showed no difference in inflammation or fungal lesions between untreated *akuB*^KU80^, Δ*aspB*, and Δ*aspB::aspB* strains [24]. This suggests the caspofungin treatment reduces growth and potentially has a fungicidal effect on the Δ*aspB* strain *in vivo*.

**Figure 4.**
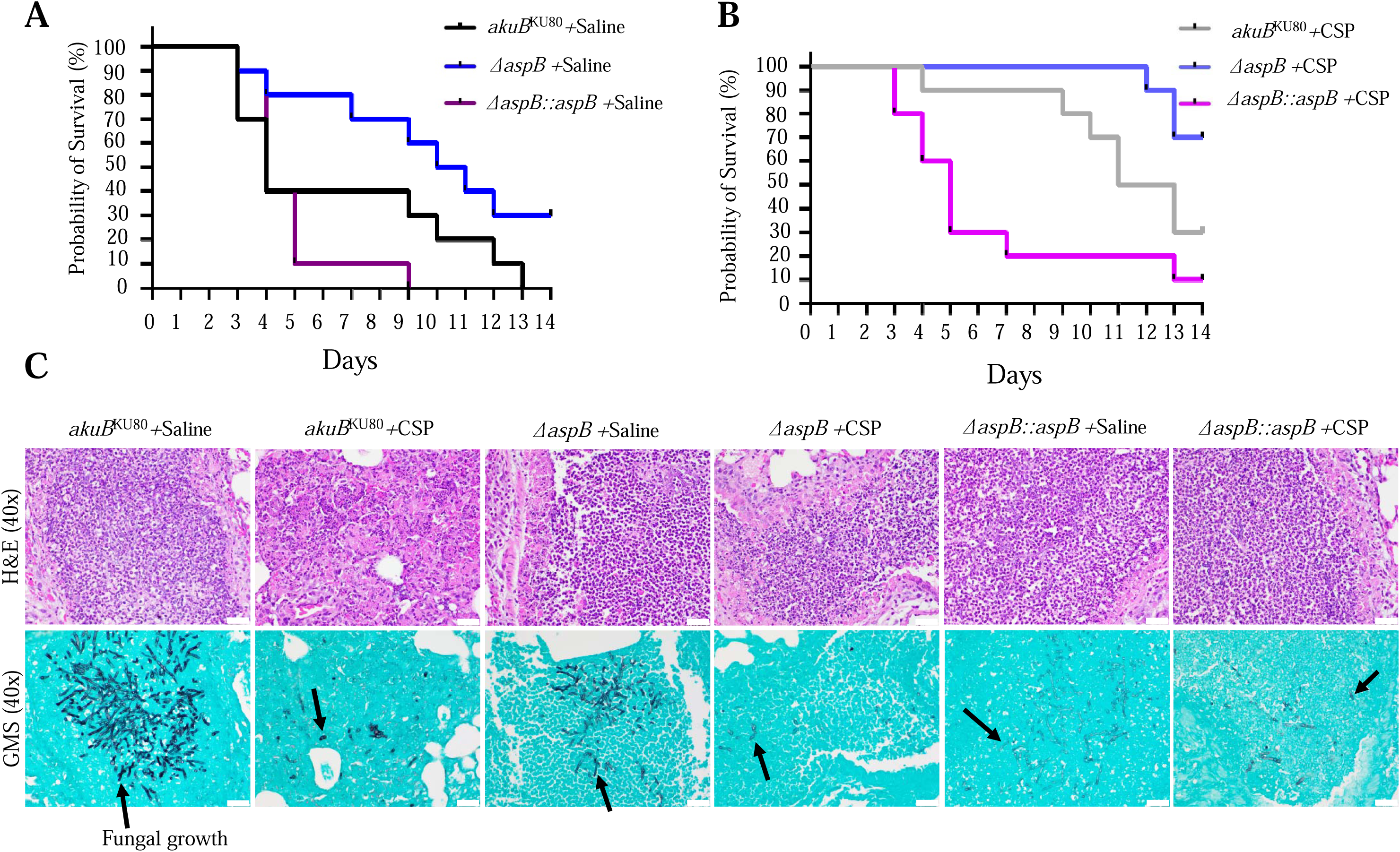
The. Δ*aspB* strain treated with caspofungin has reduced fungal burden in an immunosuppressed intranasal murine model of invasive aspergillosis. (A-B) Mice infected with Δ*aspB* strain and treated with caspofungin had a higher chance of survival. Ten mice per strain per condition were infected intranasally with 4×10^6^ conidia. Mice were treated with either saline (A) or caspofungin (B) on days +1 through +4 post-infection. Mice were monitored at least twice a day for 14 days. Survival is on a Kaplan-Meier curve with log rank pair-wise comparison (p<0.0001). (C) H&E stain of lungs after 3 days post-infection shows that there is a similar inflammatory response across conditions in the region where hyphae are present. GMS stain shows fewer and smaller fungal lesions in Δ*aspB*-infected mice treated with caspofungin. Images taken on an Echo Rebel Hybrid Light microscope using a 40x objective. Scale bar is 20 µm. Sections are 4 µm thick. Arrows denote examples of fungal growth.

### AspB Interactome Changes Post Caspofungin Exposure

Previous work determined that septins’ localization is altered by exposure to caspofungin in *C. albicans* and *A. fumigatus* [24,29,30]. Additionally, the protein interactome of septin AspB was altered after exposure to caspofungin in a qualitative proteomic experiment [32]. This approach only detected for the presence or absence of interactant proteins between basal and caspofungin conditions, potentially missing interactions that occur in both conditions but change in abundance [32]. To gain a more mechanistic insight into how AspB contributes to the fungal response to caspofungin, we applied affinity purification coupled with quantitative proteomics.

An AspB-eGFP expressing strain was grown in GMM and GMM supplemented with 1 μg/mL caspofungin for 24 hours. AspB-GFP was then purified using a GFP-Trap® affinity matrix, and proteins bound to AspB-GFP were prepared and analyzed by LC-MS/MS to identify protein interactors in each condition. Principal Component Analysis (PCA) indicates that the GMM- and caspofungin-AspB interactome are distinctive from each other (Fig. S1A). A total of 226 proteins were significantly decreased (fold change (FC) < -2, p < 0.05) and 106 proteins were significantly increased (FC > 2, p < 0.05) upon caspofungin treatment (Fig. S1B). Among the proteins increased upon caspofungin exposure was PpoA, a characterized fatty acid monooxygenase. PpoA was also identified by a previous study investigating the AspB interactome post-caspofungin exposure, indicating that our analyses were able to confirm previously described caspofungin-specific AspB interactions [32]. Candidate protein interactors that met the screening criteria (FC > 2 or FC<-2, p < 0.05) were analyzed using FungiFun3, a gene ontology tool, to assess changes in biological processes [33]. Proteins that are known to be involved in hyphal growth and cell wall-related processes were overrepresented among the list of proteins increased upon caspofungin treatment (Fig. S1C). In contrast, proteins that are known to be involved in protein folding were overrepresented in the list of proteins decreased upon caspofungin treatment (Fig. S1D).

### Δ**bgt1,** Δ**gel2,** ΔnsdD, and ΔmapA Have a Reduction in Conidiation, While ΔnsdD Has a Growth Defect

Based on our proteomics and gene ontology analyses, we narrowed down to six genes that met the following criteria for further investigation: were significantly increased after caspofungin exposure (p < 0.05), were at least two-fold greater in abundance (FC > 2), and had either a known or putative role in cell-wall related functions (Table S3). We performed the individual deletions of all six candidate genes (Table S3) to determine their role in septin-related phenotypes. We first characterized each deletion strain to understand how the gene affects growth during basal conditions. Due to the Δ*punA* not being viable at 37 °C, we decided not to pursue it further in this report. Major and biologically relevant radial growth defects are seen in Δ*nsdD* (37.8 mm average diameter compared to the 85 mm diameter of the *akuB*^KU80^ strain)(Fig. 5A). The slight reduction of radial growth in Δ*aspB* was also seen in a previous study and deemed unlikely to be biologically relevant in basal conditions, which is also the case for the Δ*gel2* strain [24]. Similarly, a previously noted defect in conidiation was seen in Δ*aspB* [24]. Δ*bgt1,* Δ*gel2,* Δ*nsdD*, and Δ*mapA* also have reduced conidiation, but not as severely as Δ*aspB* (Fig. 5B). As prior work noted that Δ*aspB* strain had delayed septation, we were interested in determining if any of the candidate gene deletion strains were also defective in septation [24].

**Figure 5.**
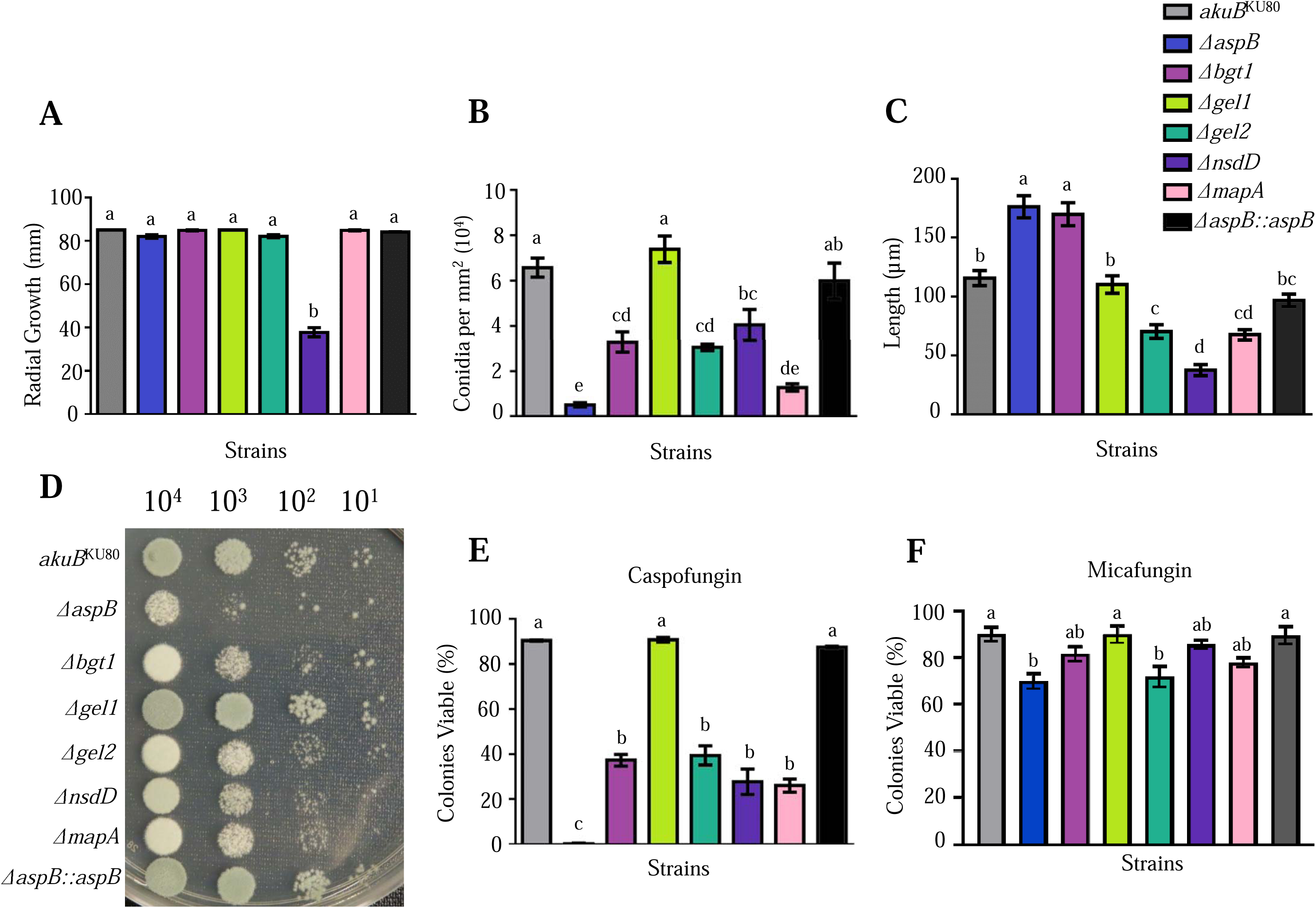
Deletion of *bgt1*, *gel2*, *nsdD*, and *mapA* increases susceptibility to caspofungin. (A) Δ*nsdD* has a significant defect in radial growth after 5 days incubation. 10^4^ total conidia of each strain were placed on GMM agar. Plates were incubated for 5 days and radial growth was measured every 24 hours. Experiments were replicated three times. (B) Deletion of *aspB, bgt1, gel2, nsdD*, or *mapA* results in reduced conidial production. Conidia were collected on day 5 of growth on GMM media using 10 mL of 0.05% Tween-80. Conidia were counted using a hemocytometer, and values were divided by the total area of growth to account for strains with growth defects. (C) Δ*aspB* and Δ*bgt1* have an increase in the length of the apical compartment. Δ*gel2,* Δ*nsdD*, and Δ*mapA* have a decrease in the length of the apical compartment. 10^4^ total conidia of each strain were inoculated onto coverslips immersed in GMM and incubated for 15 hours. Coverslips were stained with aniline blue and visualized. The length of the apical compartments (N=20) was measured using imageJ. (D) Spore dilution assays show Δ*bgt1*, Δ*gel2*, Δ*nsdD*, and Δ*mapA* strains have an increase in susceptibility to caspofungin. Conidia (10^4^-10^1^) were grown on GMM media supplemented with 1 μg/mL caspofungin for 48 hours. (E, F) Deletion of *bgt1*, *gel2*, *nsdD*, and *mapA* results in reduced, but not loss of, viability when grown in caspofungin but not micafungin. Deletion of *gel1* does not affect viability when exposed to caspofungin or micafungin. 10^4^ conidia were grown in GMM supplemented with either 1 µg/mL (E) caspofungin or (F) micafungin for 48 hours, then incubated in CFDA and visualized. Error bars represent SEM. Experiments were replicated three times. One-way ANOVA with Tukey’s multiple-comparison test was performed in Graphpad Prism and was declared significantly different at a p-value of <0.05. Group means with different lowercase letters are significantly different.

To test this, we measured apical compartment length as an indirect method of measuring potential septation defects. Δ*bgt1* mutants have a similar increase in apical compartment length to Δ*aspB*, suggesting a delay in septation (Fig. 5C). In contrast, Δ*gel1* mutants have apical compartments similar to the *akuB*^KU80^ wild-type strain (Fig. 5C), suggesting that Gel1 is dispensable for septa formation. Δ*gel2* (70.4 µm) and Δ*mapA* (67.7 µm) mutants exhibit hyperseptation. Δ*nsdD* (37.7 µm) has a more drastic reduction in apical compartment length, but this phenotype may be due to its growth defect (Fig. 5C)[34].

### Δ**bgt1,** Δ**gel2,** ΔnsdD, and ΔmapA Have Increased Sensitivity to Caspofungin Exposure

Since the candidate proteins showed increased interaction with AspB during caspofungin exposure, we explored their potential role in mediating the AspB-dependent fungal response to caspofungin. To test this, we conducted a spore dilution assay on GMM agar supplemented with 1 μg/mL caspofungin. 10^4^, 10^3^, 10^2^, and 10^1^ spores were plated and incubated at 37°C for 2 days. Deletion of the *aspB* gene displayed colony-level growth defects in the 10^3^, 10^2^, and 10^1^ concentrations (Fig. 5D). No other deletion strains displayed as severe of a sensitivity to caspofungin. Δ*bgt1,* Δ*gel2,* Δ*nsdD*, and Δ*mapA* lost full colony growth at 10^2^ and 10^1^ concentrations (Fig. 5D). In contrast, Δ*gel1* is similar to the *akuB*^KU80^ wild-type and Δ*aspB::aspB* complemented strains (Fig. 5D). The minimum effective concentration (MEC) of caspofungin for Δ*bgt1,* Δ*gel2,* Δ*nsdD*, and Δ*mapA* was determined to be lower than that of the wild-type, and equal to or lower than that of Δ*aspA*, Δ*aspB,* and Δ*aspC* (Table S4).

### Reduced Viability of Δbgt1, Δgel2, ΔnsdD, and ΔmapA Post-Caspofungin Exposure

Since *aspB* deletion leads to a fungicidal response to caspofungin, we investigated whether any candidate gene deletion strains also exhibited reduced viability. All strains showed equal viability in basal conditions, confirmed qualitatively by visible fluorescence present throughout the mycelium (Fig. 6A). We then grew all strains for 48 hours in the presence of 1 µg/mL caspofungin and determined their viability with CFDA. The *aspB* deletion strain showed nearly no viability at 37°C, as previously shown in Figure 1B (Fig. 5E, Fig. 6B). No other strain demonstrated the complete loss of viability phenotype observed in the Δ*aspB*. The Δ*bgt1*, Δ*gel2*, Δ*nsdD*, and Δ*mapA* strains form microcolonies in the presence of caspofungin like the *akuB*^KU80^ strain, but they had a reduction in viability (Δ*bgt1*=37.4%, Δ*gel2*=39.5%, Δ*nsdD*=27.8%, Δ*mapA*=26.1% compared to *akuB*^KU80^ =90.4%)(Fig 5E). Δ*gel1* did not have a reduction in cell viability when grown in caspofungin (Δ*gel1*=90.8%) (Fig. 5E, Fig. 6B). To determine if any of the candidate gene deletion strains had a unique phenotype to other echinocandins, we conducted the same assay but with micafungin, similar to Figure 1C. The candidate gene deletion strains do not have a high reduction of viability when grown in micafungin (Fig. 5F, Fig. S2).

**Figure 6.**
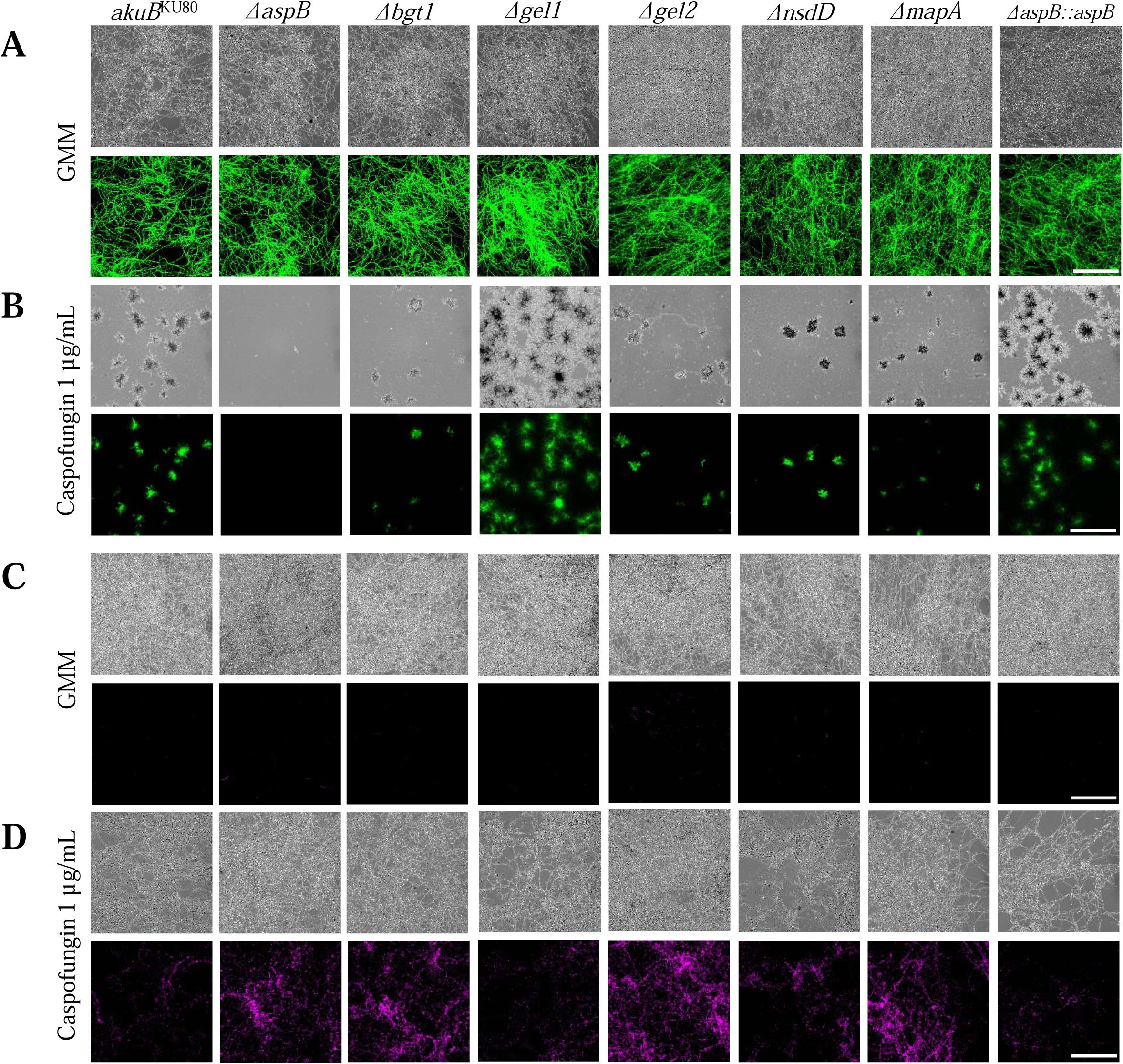
Δ***bgt1*,** Δ***gel2*,** Δ***nsdD*, and** Δ***mapA* strains have reduced cell viability and increased hyphal damage from caspofungin.** (A) All strains were fully viable when grown in GMM media. Conidia (10^4^) were grown in GMM for 24 hours on coverslips then incubated in CFDA. Slides were prepared and visualized on an Inverted Leica DMi8 with Leica K5 Microscope Camera using a 10x objective. (B) Candidate deletion strains did not lose total viability, but strains Δ*bgt1*, Δ*gel2*, Δ*nsdD*, and Δ*mapA* had a partial reduction in viability. Conidia (10^4^) were grown on coverslips immersed in GMM supplemented with 1 μg/mL caspofungin for 48 hours. Coverslips were then incubated in CFDA and visualized. (C) Hyphal damage was not observed in basal conditions. Conidia (10^4^) were grown on coverslips in GMM for 24 hours. Coverslips were washed in PIPES (pH 6.7) for 5 minutes twice then treated with propidium iodide (PI) solution. Coverslips were washed twice with PIPES, then visualized on an Inverted Leica DMi8 with Leica K5 Microscope Camera using a 10x objective. (D) Strains Δ*bgt1*, Δ*gel2*, Δ*nsdD*, and Δ*mapA* had hyphal damage when treated with caspofungin. Conidia (10^4^) were grown on coverslips in GMM for 24 hours, then incubated in caspofungin for 2 hours. Cells were then washed with PIPES and incubated with PI. Experiments were performed at 37°C. All experiments were replicated three times. Scale bar is 500 µm.

### Bgt1, Gel2, NsdD, and MapA are Involved in the Response to Caspofungin in Mature Mycelium

Next, we determined if any of our candidate gene deletion strains were susceptible to hyphal damage when exposed to caspofungin as mature mycelia. To visualize hyphal damage, we utilized a propidium iodide (PI) stain. No hyphal damage was observed in basal conditions, indicating that there is no defect in the cell wall at basal conditions (Fig. 6C). The experiment was then repeated with a 2-hour incubation in GMM supplemented with 1 μg/mL caspofungin prior to PI staining. Extensive hyphal damage was seen in Δ*bgt1*, Δ*gel2*, Δ*nsdD*, and Δ*mapA* strains, similar to Δ*aspB,* after exposure to caspofungin (Fig. 6D). Taken together, Bgt1, Gel2, NsdD, and MapA are involved in the fungal response to caspofungin in earlier stages of growth and mature hyphae.

### Candidate Genes are not involved in the response to other Cell Wall Disrupting Agents or in the Caspofungin Paradoxical Effect

We further characterized the candidate gene deletion strains’ sensitivity to other cell wall-disturbing agents. To test this, we plated conidia on GMM agar and GMM supplemented with either 1 μg/mL caspofungin, 100 μg/mL Congo red, 2 μg/mL nikkomycin Z, 5 μg/mL calcofluor white, or 10 μg/mL calcofluor white and incubated for three days. Additionally, we plated 4 μg/mL caspofungin and incubated for five days to observe whether the deletion strains were still capable of the caspofungin paradoxical effect. This paradoxical effect, sometimes known as the Eagle phenomenon, is when increasing concentrations of a drug become less effective. After incubation, Δ*aspB* shows increased susceptibility to caspofungin and Congo red, and slight increase in susceptibility to nikkomycin Z and calcofluor white (Fig. S3). We observed Δ*gel1* had a mild increase in susceptibility to Congo red compared to the *akuB*^KU80^ wild-type (Fig. S3).

Δ*bgt1*, Δ*gel2*, and Δ*nsdD* had a mild increase in susceptibility to 10 μg/mL calcofluor white, but not as pronounced as Δ*aspB* (Fig. S3). To avoid missing potential phenotypes, we also conducted spot dilution assays with these drugs (Fig. S4). We again observed a slight increase in susceptibility of Δ*bgt1*, Δ*gel2*, and Δ*nsdD* to calcofluor white, but no novel phenotypes were seen (Fig. S4). Taken together, these candidate genes might mediate fungal response to caspofungin while not being a part of a general fungal cell wall stress response.

### Candidate genes are involved in the fungal response to caspofungin through the same mechanisms as AspB

To further confirm that AspB is functioning in the same pathways as the candidate proteins, we generated double deletion mutants that lack both the *aspB* and one of the genes that previously had a phenotype to caspofungin (*bgt1*, *gel2*, *nsdD*, and *mapA*). As these strains appeared slightly more sickly, we decided to verify that they do not have a phenotype in basal conditions that could largely affect the interpretation of drug assays. We conducted radial growth, conidial quantification, and spore viability assays. The Δ*aspB*Δ*nsdD* strain had a similar growth defect to Δ*nsdD* (Fig. 7A). The other double mutants have typical radial growth, resembling either Δ*aspB* or their other parent (Fig. 7A). All double mutant strains had reduced conidia production, similar to their Δ*aspB* parent, or both Δ*aspB* and Δ*mapA* in the case of the Δ*aspB*Δ*mapA* strain (Fig. 7B). Spores were tested for viability by staining 1x10^7^ conidia of each strain with propidium iodide for inviability and calcofluor white as confirmation of being a spore. Heat-killed wild-type spores were used to standardize images. All strains had viable spores, albeit all mutants had slightly more inviable spores than the wild-type (Fig. 7C). To test for mycelial viability, a CFDA assay was conducted, similar to that of Figures 2 and 6. All strains were viable under basal conditions. Strains were then grown for 48 hours in the presence of 1 µg/mL caspofungin and we determined their viability with CFDA. All double deletion strains had similar viability reduction as the *aspB* deletion strain (Fig. 7D). To test if there is increased hyphal damage in mature mycelium, we used propidium iodide staining on mycelium grown for 24 hours. Little to no fluorescence was observed, suggesting no major defect in the cell wall under basal conditions (Fig. S5A). This experiment was repeated with a 2-hour incubation in GMM supplemented with 1 μg/mL caspofungin, followed by PI staining. Similar to both Δ*aspB* and their other parent, all double deletion strains had increased hyphal damage post-caspofungin exposure (Fig. S5B).

**Figure 7.**
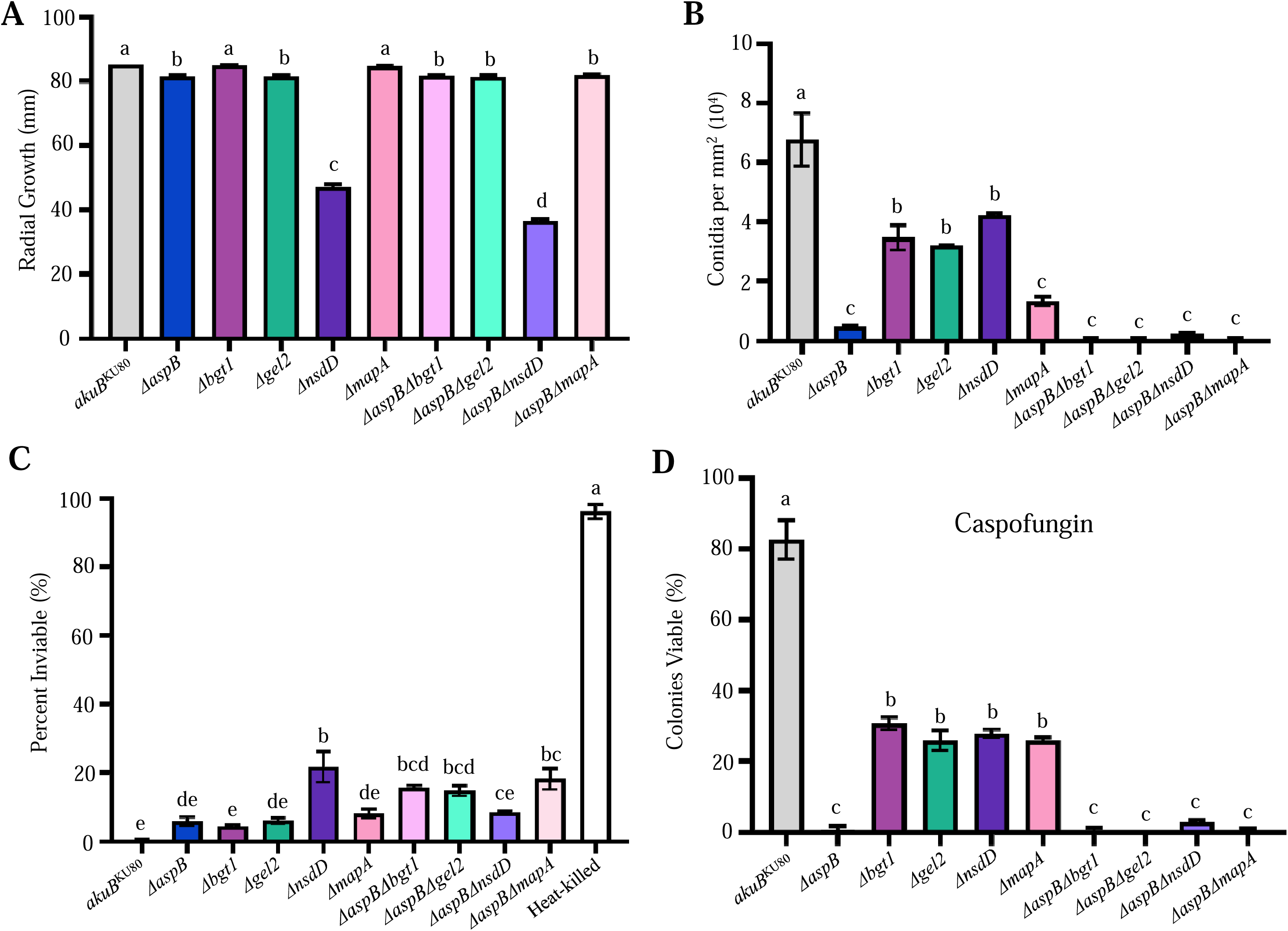
Double-deletion mutants are similar to the. Δ***aspB* single mutant under caspofungin exposure.** (A) Δ*aspB*Δ*nsdD* radial growth defect is similar to that of the Δ*nsdD* parent. 10^4^ total conidia of each strain were placed on GMM agar. Plates were incubated for 5 days and radial growth was measured every 24 hours. (B) Double deletion mutants resemble Δ*aspB* conidiation defect. Conidia were collected on day 5 using 10 mL of 0.05% Tween-80. Conidia were counted via hemocytometer and values were divided by the total area of growth to account for strains with growth defects. (C) Deletion mutants’ conidia are viable. 10^7^ spores were spun down, washed with PIPES twice, and incubated with propidium iodide and calcofluor white for 5 minutes. Spores were washed twice again, then imaged and counted for viability. (D) Δ*aspB* and double mutants have fungicidal response to caspofungin. 10^4^ conidia were grown in GMM supplemented with 1 µg/mL caspofungin for 48 hours then incubated in CFDA and visualized. Viable colonies were counted. All experiments done in triplicate. Error bars represent SEM. One-way ANOVA with Tukey’s multiple-comparison test were performed in Graphpad Prism and were declared significantly different at a p-value of <0.05. Group means with different lowercase letters are significantly different.

Taken together, these data suggest that AspB contributes to the candidate proteins role in response to caspofungin.

## Discussion

The septins are a highly conserved family of cytoskeletal proteins with a variety of cellular roles. This includes roles in cell division, stress response, cytoskeleton organization, and scaffolding [24,35–37]. Previous work noted the role of septins in response to cell wall stress. In *A. fumigatus,* it was determined that the septins have a role in response to cell wall stress, notably showing that Δ*aspB* exhibits a hypersensitive response to caspofungin exposure on solid media [24]. Here, we aimed to further investigate the role of septins in the *A. fumigatus* response to caspofungin. We found that deletion of *aspA, aspB*, and *aspC* reduced fungal viability after caspofungin exposure and increases hyphal damage in mature mycelia, whereas deletion of *aspB* elicited a fungicidal response. The drastic phenotype of the Δ*aspB* strain was surprising, as *Aspergillus spp.* septins form heteropolymers consisting of hexamers (AspA-C) and octamers (AspA-D) [38]. Thus, it would be expected that deletion of genes that abolish septin heteropolymers would have similar phenotype. Deletion of *aspD* still allows proper formation of hexamers but not octamers. In our work, we found that the *aspD* deletion strain was slightly more sensitive than the *akuB*^KU80^ wild-type, Δ*aspB::aspB*, or Δ*aspE* strains, although its increased susceptibility was not as pronounced as that of the *aspA*, *aspB*, and *aspC* deletion strains (Fig. 1). This suggests that the presence of the hexamer is at least sufficient, if not necessary, for proper fungal response to caspofungin. It is unknown if the sole presence of the octamer would also be sufficient, as deletion of *aspA-C* prevents the formation of both structures. Similarly, Δ*aspB*’s fungicidal response to clinically relevant doses of caspofungin could be due to AspB being specifically needed to scaffold necessary proteins in response to caspofungin, regardless of the septin complex, while the Δ*aspA* or Δ*aspC* leads to a loss of both the hexamer and octamer, which could causes general dysfunction in AspB’s role in coordinating the fungal response to caspofungin. Proteomics analysis in *Cryptococcus neoformans* also supports the possibility of subunit-specific protein interactions, where the *cdc3^aspB^* and *cdc10^aspD^* septins are shown to have differences in interactomes after exposure to cellular stressors [39]. All of the septin deletion strains retain the caspofungin paradoxical effect, suggesting that individual septins and their heteropolymers act independently of the pathways that contribute to the caspofungin paradoxical effect.

Septins are involved in septation, as the Δ*aspA,* Δ*aspB,* Δ*aspC,* and Δ*aspE* strains have fewer septa and a larger apical compartment compared to the *akuB*^KU80^ strain [24]. It has been shown that septation is a critical cellular process for maintaining the fungistatic effect of echinocandins, as the deletion of genes involved in the septation initiation network (SIN) results in a fungicidal effect [31]. The Δ*aspE* strain has a septation defect of similar magnitude to the Δ*aspB* strain, but the Δ*aspE* strain does not have a drastic increase in susceptibility to the echinocandins.

Additionally, Δ*aspA* and Δ*aspC* have a septation defect, but their decrease in viability post-caspofungin exposure is not as severe as in the Δ*aspB* strains. Furthermore, deletion of SIN genes confers fungicidal activity to both micafungin and caspofungin. Thus, the septation defect could partially contribute to the increased susceptibility to echinocandins in the Δ*aspA,* Δ*aspB,* and Δ*aspC* strains; however, the septation phenotype alone is not responsible for the fungicidal effect observed in the Δ*aspB* strain during caspofungin exposure.

The echinocandin class includes four clinically approved drugs: anidulafungin, micafungin, caspofungin, and most recently approved, rezafungin [40,41]. These drugs all work through noncompetitive inhibition of the β-1,3-D-glucan synthase, but alterations in their side chains comprise the major differences between each one [18,42]. These changes in the side chains can lead to changes in how the drug impacts fungal physiology. For instance, caspofungin is the only echinocandin to induce the paradoxical effect [42]. Unlike micafungin, high concentrations of caspofungin increase the levels of cytosolic calcium [43,44]. This activates calmodulin-calcineurin signaling and leads to the caspofungin paradoxical effect [44]. Thus, our findings add to the growing evidence that even though the different echinocandins have a shared mechanism of action, how the fungus responds to echinocandins is drug-specific.

Animals infected with the Δ*aspB* strain and treated with caspofungin had a 70% chance of survival, which is higher than that of animals infected with *akuB*^KU80^ strain (30%). The first death of Δ*aspB* strain infected mice treated with caspofungin was recorded on day +12 post infection, compared to Δ*aspB* strain infected mice treated with saline or the other strains and conditions with first deaths on day +3 or day +4. It is possible that colonies that persist under caspofungin treatment are capable of growing and establishing infection after the drug pressure is released.

Additionally, while Δ*aspB* strain infected mice treated with saline still had 30% survival, this is not statistically significant. Previous work also determined loss of AspB does not impact fungal virulence and resulted in a similar murine survival rate [24]. Our histological analyses showed reduced inflammation and reduced fungal lesions in the lungs of Δ*aspB* infected mice treated with caspofungin. However, this assay is done on day +3 after infection when mice are still being treated with caspofungin. Nonetheless, this increase in survival makes AspB a prospective target for drug therapy developments, as use in conjunction with caspofungin elicits a fungicidal response in *A. fumigatus*. Currently, only the plant cytokinin forchlorfenuron (FCF) is known to disrupt septin organization [45]. This compound disrupts septins within both fungal and mammalian cells by interfering with their ability to bind and hydrolyze GTP [45–47]. It is possible that FCF could be modified and refined to target fungal-specific septins. FCF analogues have previously been developed to better target ovarian and endometrial cancers [48]. Thus, FCF analogues could be used in conjunction with caspofungin to create a new fungicidal therapy for the treatment of IA. Further work is required to better understand the mechanisms of AspB’s response to caspofungin and to develop novel clinical therapies.

Our proteomic analysis identified four candidate protein interactors of septin AspB that were significantly increased by caspofungin treatment and have a potential role in cell wall function. One limitation of our experiment is the lack of an input control, such as GFP ectopically expressed. Because of the lack of a GFP-only control, some hits may be nonspecific GFP-interactants. We verified that our candidate genes were not increased in expression after caspofungin exposure based on publicly available datasets [49]. *bgt1*, *gel2*, *nsdD*, and *mapA* all decrease in transcript abundance after caspofungin exposure, while *gel1* and *punA* increase in transcript abundance after caspofungin exposure [49]. This suggests that the increase in *bgt1*, *gel2*, *nsdD*, and *mapA* interactions observed in our AspB pulldown experiment might not be due to an increase in overall abundance. Additionally, to mitigate the possible false positives that could be obtained from this process, we performed epistasis analyses and generated double mutants of our candidate and *aspB,* in which candidate genes were epistatic to *aspB*.

Bgt1, Gel1, and Gel2 are glucanosyltransferases in *A. fumigatus* [50]. These glucanosyltransferases work to remodel unorganized β-1,3-glucan chains in the periplasmic space to stabilize and modify the cell wall as needed [50]. Due to their role in cell wall homeostasis, previous work identified these three glucanosyltransferases as potential drug targets that required further investigation [50]. Bgt1 works by hydrolyzing a β-1,3-linked oligosaccharide and placing it on another molecule of β-1,3-glucan [51]. In *C. albicans*, deletion of *BGL2*, the *bgt1* homologue, resulted in an increase in susceptibility to nikkomycin Z, a chitin synthase inhibitor [52]. In contrast, *A. fumigatus* Δ*bgt1* only has a slight increase in susceptibility to nikkomycin Z (Fig. S3). Deletion of *BGL2* in *S. cerevisiae* increased the cellular chitin content [53]. In *A. fumigatus,* though, Δ*bgt1*Δ*bgt2* double mutants do not have a change in chitin content [51]. A slight reduction in conidiation was observed in *A. fumigatus* Δ*bgt1* (Fig. 5A, 5B). *A. fumigatus* Δ*bgt1* susceptibility to calcofluor white and Congo red were similar to the *akuB*^KU80^ wild-type, although compared to basal conditions there was a reduction in growth (Fig. S3) [54]. A potential mechanism for the increased susceptibility in *A. fumigatus* Δ*bgt1* and Δ*aspB* to caspofungin is that hyphal damage may be exacerbated by the lack of interaction between AspB and Bgt1 when either party is missing (Fig. 5E). Due to the role of septins in septation and scaffolding, the loss of AspB in particular may be more deleterious than just the loss of Bgt1 and thus result in the observed fungicidal effect (Fig. 1B). Gel1 and Gel2 are two GPI-anchored glucanosyltransferase members of the Glycoside Hydrolase Family 72 (GH72) [55]. Despite having the same enzymatic activity, deletion of *gel1* did not result in a unique phenotype, but reduced mycelial growth and abnormal cell wall architecture was observed in Δ*gel2* during basal conditions (Fig. 5A-B) [50,51,56]. An increase in chitin is associated with deletion of *gel2*, a compensatory mechanism similar to deletion of *bgt1* [56]. *A. fumigatus* Gel1 is dispensable for proper fungal response to caspofungin (Fig. 5D-E). In contrast, Gel2 does appear to be involved in the response to caspofungin due to increased susceptibility to caspofungin and reduced viability in caspofungin compared to *akuB*^KU80^ wild-type (Fig. 5D-E)[56]. Our proteomics analysis noted an increased interaction between Gel2 and AspB, but cellular levels of Gel2 were not explored. It is unknown if cellular levels of Gel2 are increased in *A. fumigatus* post-caspofungin exposure, similar to *C. albicans*. Additionally, further work exploring the overexpression of Gel2 may uncover a new mechanism of resistance to caspofungin in *A. fumigatus*. Further work is also needed to uncover the relationship between glucanosyltransferase and the septins in septation and cell wall stress responses.

NsdD (**<u>n</u>**ever in **<u>s</u>**exual **<u>d</u>**evelopment) is a putative GATA-type transcriptional activator with roles in *A. nidulans* for sexual development [57]. NsdD is responsible for activating sexual development [58]. Deletion of *nsdD* results in the inability to produce cleistothecia (fruiting bodies), while overexpression allows production of Hülle cells even in conditions which typically block sexual development [58]. Despite being a primarily asexual fungus, *A. fumigatus* is capable of and still holds a variety of functional sexual reproductive genes [59]. NsdD has been implicated in cell wall remodeling and hyphal fusion [60]. Deletion of *nsdD* results in reduced hyphal growth on minimal media (Fig. 5B) [57,60]. Drug challenges show some sensitivity of ΔnsdD to Congo red [60] and caspofungin (Fig. 5D-E, Fig. S3). Previous work also noted weakened hyphal tips in Δ*nsdD*, which is in accordance with the increased hyphal damage during cell wall stress seen via PI staining (Fig. 6D)[60]. We suspect that while NsdD may have independent roles in sexual development, its previous cell wall remodeling functions during hyphal fusion for heterokaryon formation may have alternative roles in *A. fumigatus* cell wall remodeling outside of sexual development. Further work is required to understand the NsdD-mediated transcriptional response to caspofungin exposure.

MapA (gene *Afu4g06930*) is a previously uncharacterized protein in *A. fumigatus*. In *S. cerevisiae* and *C. albicans*, cytosolic protein MAP2 is an ortholog that functions as a methionine aminopeptidase. Deletion of MAP2 results in slightly slower growth rate and some chemical sensitivities in *S. cerevisiae* and *C. albicans* [61,62]. It is possible that this methionine aminopeptidase can remove the methionine to promote the maturation of proteins involved in the response to caspofungin. However, which proteins are regulated by MapA in *A. fumigatus* is unknown.

Although septins have been implicated in the mediation of cell wall stress, the full mechanism how they contribute to this phenotype is not fully understood [23,24,28,29,32]. Previous proteomic analyses and our own proteomics experiments show that MpkA, MkkA, and other cell wall integrity components co-immunoprecipitated with AspB [32]. It is possible that AspB interacts with the cell wall integrity pathway to facilitate fungal response to caspofungin [32]. In *A. nidulans,* deletion of *aspB* partially rescues both the growth defect and susceptibility to caspofungin of the Δ*mpkA* strain, further hinting at a possible crosstalk between the septin and the CWI [23]. As we did not observe a difference in abundance in the CWI pathways kinases between basal and caspofungin conditions, we did not explore the possible role of septins in regulating the CWI. Nonetheless, further work will be needed to investigate this possibility. As we see different pathways differentially interacting with AspB, we hypothesize that the fungicidal response observed in the Δ*aspB* strain is not due to a singular AspB-mediated interaction that is yet to be found, but instead by AspB having a global effect on a number of contributing proteins and pathways. Future studies directed towards developing drugs specific to fungal septins or in determining additional septin interactants that have key roles in the caspofungin response can help to better understand ways in which caspofungin therapy can be improved for use against invasive aspergillosis.

## Materials and Methods

### Strains, media, and conditions

*A. fumigatus akuB*^KU80^ served as the wild-type and control strain. Δ*aspB::aspB* complemented strain served as an additional control to determine that phenotypes were due to the deletion of the *aspB* gene. Septin deletion strains are described in (Table S1, [24,63]). The *aspB-egfp* strain from [24], with *aspB-egfp* expressed by the *aspB* promoter from the native loci, was used to pulldown AspB interactants. All cultures were plated on glucose minimal media (GMM) supplemented with 5 mM uracil and 5 mM uridine (GMM+UU) and incubated at 37°C, unless otherwise specified.

### Drug susceptibility assays

Spores were counted, diluted, and inoculated in rows onto agar plates containing caspofungin (1 μg/mL) in 10μL quantities with a total of 10^4^, 10^3^, 10^2^, and 10^1^ conidia. Plates were incubated for 48 hours at 37°C unless otherwise noted. For the minimum effective concentration assay, a quantity of 2.5x10^4^ spores were added to RPMI media with different concentrations of caspofungin and incubated for 48 hours at 37°C according to CLSI guidelines [64]. We were not able to test the *aspD* and *punA* deletion strains according to CLSI guidelines as Δ*aspD*, which is an uracil/uridine auxotroph, could not grow in RPMI, and Δ*punA*, which has a temperature sensitivity, could not grow at 37°C.

### Fungal Viability Assay

Conidia of *akuB*^KU80^, Δ*aspA-E*, and Δ*aspB::aspB* were diluted to 10^4^ spores and cultured on coverslips immersed in 4 mL of GMM+UU broth and GMM+UU+caspofungin (1 μg/ml) and incubated at 37°C for 24 and 48 hours, respectively. To examine cell viability, coverslips were incubated in 5-carboxyfluorescein diacetate (CFDA) (50 μg/mL 0.1 M MOPS pH 3) for 1 hour at 37°C and 70 rpm [31]. Slides were prepared for microscopy and image on an Inverted Leica DMi8 with Leica K5 Microscope Camera using a 10x objective. Images were analyzed using imageJ [65]. Viability in basal conditions (GMM) was determined qualitatively by noting the presence of visible fluorescence throughout the mycelium. To quantitatively determine viability when grown in caspofungin, viable, fluorescent colonies were manually counted and divided over total colonies counted. A minimum of 50 colonies per strain per replicate were counted.

### Spore Inviability Assay

Conidia of *akuB*^KU80^, heat-killed *akuB*^KU80^ (5 minutes at 95°C), Δ*aspB*, and the single and double deletion mutants were diluted to 10^7^ spores and spun down via centrifuge for 5 minutes at maximum speed. Their media was removed and replaced with PIPES (50 mM PIPES, pH 6.7), then vortexed to dislodge the pellet. Spores were washed a total of two times in this manner, then vortexed and incubated with calcofluor white (1 µg/mL) and propidium iodide (12.5 µg/mL) for 5 minutes at room temperature. Spores were spun down and washed with PIPES two more times. Slides were prepared for microscopy and imaged on Nikon Ti-Eclipse microscope using a 100x objective. Images were analyzed using imageJ [65]. Inviability was determined by standardizing against the heat-killed sample, then manually counted for the number of inviable spores and divided over total spores counted [66]. A minimum of 50 spores per strain per replicate were counted.

### Hyphal Damage Assay

Conidia (10^4^) of *akuB*^KU80^, Δ*aspA-E*, and Δ*aspB::aspB* were cultured on coverslips immersed in 4 mL of GMM+UU broth and incubated for 24 hours at 37°C. Coverslips were then incubated with GMM+UU or GMM+UU+caspofungin (1 μg/ml) for 2 hours at 37°C. To examine the damage of mature hyphae, coverslips were washed with 4 mL PIPES (pH 6.7) for 5 minutes.

PIPES was then removed and washed again with 4 mL PIPES for 5 minutes. After removing the second PIPES wash, 500 uL of propidium iodide (PI) solution (12.5 ug/mL in 50 mM PIPES) was added on top of the coverslip and incubated in the dark for 5 minutes. Slides were washed in 4 mL PIPES twice as described previously. They were then prepared for microscopy and imaged on an Inverted Leica DMi8 with Leica K5 Microscope Camera using a 10x objective. Images were analyzed in imageJ [65].

### Neutropenic Murine Model of Invasive Aspergillosis

Murine experiments followed previously established intranasal neutropenic models of IA [67,68]. Sixty 6-week old male CD1 mice (Charles River Laboratories, Raleigh, NC) were injected via intraperitoneal route with 175 mg/kg cyclophosphamide on days −2 and +3 and 40mg/kg triamcinolone acetonide subcutaneously on days −1 and + 6. Twenty mice per strain were infected intranasally with 40 μL of 10^8^ spores/mL conidia suspension of the *akuB*^KU80^, Δ*aspB*, or Δ*aspB::aspB* strains on day +0. On days +1 through +4, mice were injected via intraperitoneal route with either 2 mg/kg caspofungin or saline. Mice were monitored until day +14 and humanely euthanized via CO_2_ followed by cervical dislocation if they showed severe symptoms.

Survival was plotted on a Kaplan-Meier curve with a log rank pair-wise comparison. Murine experiments were conducted in compliance with the SIU Institutional Animal Care and Use Committee Protocol 20-034 and Virginia Tech Protocol 23-254.

### Histopathology Analysis

Mice were immunocompromised and treated as described above in [Section: *Neutropenic murine model of Invasive Aspergillosis*]. Lungs were harvested on day +3 after infection via isoflurane euthanasia and cervical dislocation. Tissue sections were stained using hematoxylin and eosin (H&E) stains to visualize inflammation and Gomori’s methenamine silver stain to visualize fungal hyphae.

### Protein Extraction, AspB-eGFP Fusion Protein Purification, and LC-MS/MS Analysis

The *aspB-egfp* strain from [24] was grown in GMM liquid media and GMM liquid media supplemented with 1 μg/mL of caspofungin for 24 hours at 37°C. Protein extraction and pulldown were completed as described by [32,69]. Fungal mycelia were homogenized to obtain total cell lysate and AspB complexes were purified using the GFP-Trap® affinity purification (Chromotek), as described [69]. GFP-Trap® magnetic beads were equilibrated by washing beads three times in 500 μL ice-cold dilution buffer then resuspended in 100 μL ice cold dilution buffer. The resin suspension is then mixed with total cell lysate (10 mg total protein) and incubated at 4°C with gentle agitation for 2 hours. Beads were collected using a magnetic stand. Beads were washed in 500 μL ice-cold lysis buffer and five times with 500 μL of wash buffer.

### Beads were suspended in 50 μL wash buffer

Samples were digested on-bead with trypsin followed by C18 desalting. Samples were analyzed via LC-MS/MS on a Thermo Easy nLC 1200-QExactive HF in technical duplicate. All mass spectra data was processed using MaxQuant (ver. 1.6.12.0) and searched against the Uniprot *Aspergillus fumigatus* proteome reviewed database (Proteome ID UP000002530). MaxQuant output was further processed via Perseus with filtering at 1% false discovery rate (FDR). Only proteins with >1 peptide were reported.

### Prioritization of Candidate Genes and Generation of Deletion Mutants

Candidate genes were chosen by prioritizing proteins with at least a two-fold increase, statistical significance, and known or putative roles in cell wall functions as listed in FungiFun3 GO term search for biological processes (Table S3) [33]. Deletions of *bgt1* (*Afu1g11460*; fungidb.org), *gel1* (*Afu2g01170*; fungidb.org), *gel2* (*Afu6g11390*; fungidb.org), *nsdD* (*Afu3g13870*; fungidb.org), *mapA* (*Afu4g06930*; fungidb.org), and *punA* (*Afu6g07470*; fungidb.org) genes were obtained by replacing the gene with the 2.4 kb *pyrG* gene from *Aspergillus parasiticus*.

Approximately 1 kb of promoter and terminator region of each gene were PCR-amplified from AF293 genomic DNA. Deletion constructs were generated by overlap fusion PCR and subsequently transformed into *akuB*^KU80^ *pyrG*− strain, all as previously described by [67].

Primers used in the transformation of all candidate strains are found in Table S2. Transformants were validated via PCR screening.

### Deletion of *aspB* via CRISPR/Cas9-Mediated Transformation

Double deletion and non-*akuB*^KU80^ Δ*aspB* strains were generated by deleting the *aspB* gene in the Δ*bgt1,* Δ*gel2,* Δ*nsdD,* Δ*mapA,* AF293, and CEA10 backgrounds. CRISPR/Cas9-mediated transformation was performed, as previously described by [70]. For the homologous repair template, a 2890 bp hygromycin B resistance (*hph*) cassette from the pUCGH plasmid was PCR amplified with 35 bp of *aspB* microhomology domains at each end. crRNAs were designed to target outside the *aspB* gene. The approximately 1.8 kb *aspB* gene was replaced with the total 2960 bp *hph* plus *aspB* microhomology domains in each of the strains. The crRNA sequences used are in Table S2. Deletion of *aspB* was verified via PCR by screening for the lack of the *aspB* gene, total size of the construct, and presence of *hph*. Primers used are in Table S2.

## Acknowledgments

R.J.B. was supported by the Southern Illinois University Carbondale Master’s Fellowship and Virginia Tech GRAD Fellowship. R.J.B. and J.V.M. were supported in part by 1R01AI165656-01A1. This work was supported in part by the College of Science at Virginia Tech (Fund BR-COS1357) and made possible by funding from the Fralin Life Sciences Institute by FLSI Award 260027. We also want to thank the members of the Vargas-Muñiz group for their critical reading of the manuscript. We want to thank Stacey Mcgee (SIU histology core facility) and Dr. Jennifer Harris (SIU Laboratory Animal Program Director) for providing guidance and support during animal experiments.

---

**Supplementary Figure 1.**
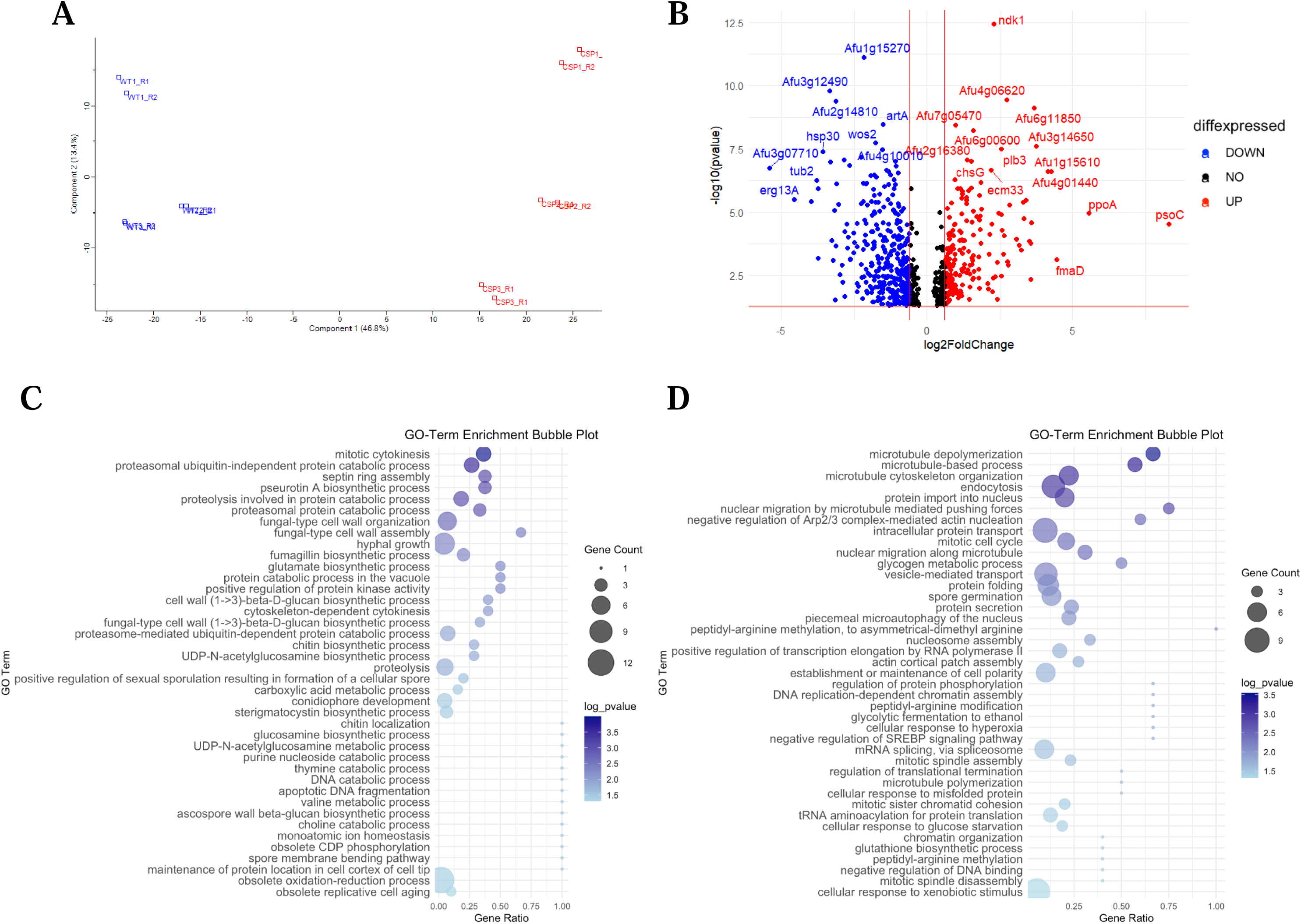
Caspofungin exposure alters AspB protein interactions. (A) Principal component analysis (PCA) of the proteomics data reveals different protein composition from AspB pulldown from GMM and caspofungin treated samples. Blue color denotes those grown in GMM, red color denotes fungi grown in GMM+ 1 μg/mL caspofungin. (B) 226 significantly decreased (FC < -2, p < 0.05) proteins (blue) and 106 significantly increased (FC > 2, p < 0.05) proteins (red). Proteins under two-fold change or had a p-value greater than 0.05 are shown in black. Red horizontal line denotes p=0.05, red vertical lines denote ± 2 fold change. (C) Category of biological processes most enriched after exposure to caspofungin is hyphal growth. (D) Category of biological processes most decreased after exposure to caspofungin is protein folding. (C,D) Gene ontology enrichment for biological processes determined by using FungiFun3 [33].

**Supplementary Figure 2.**
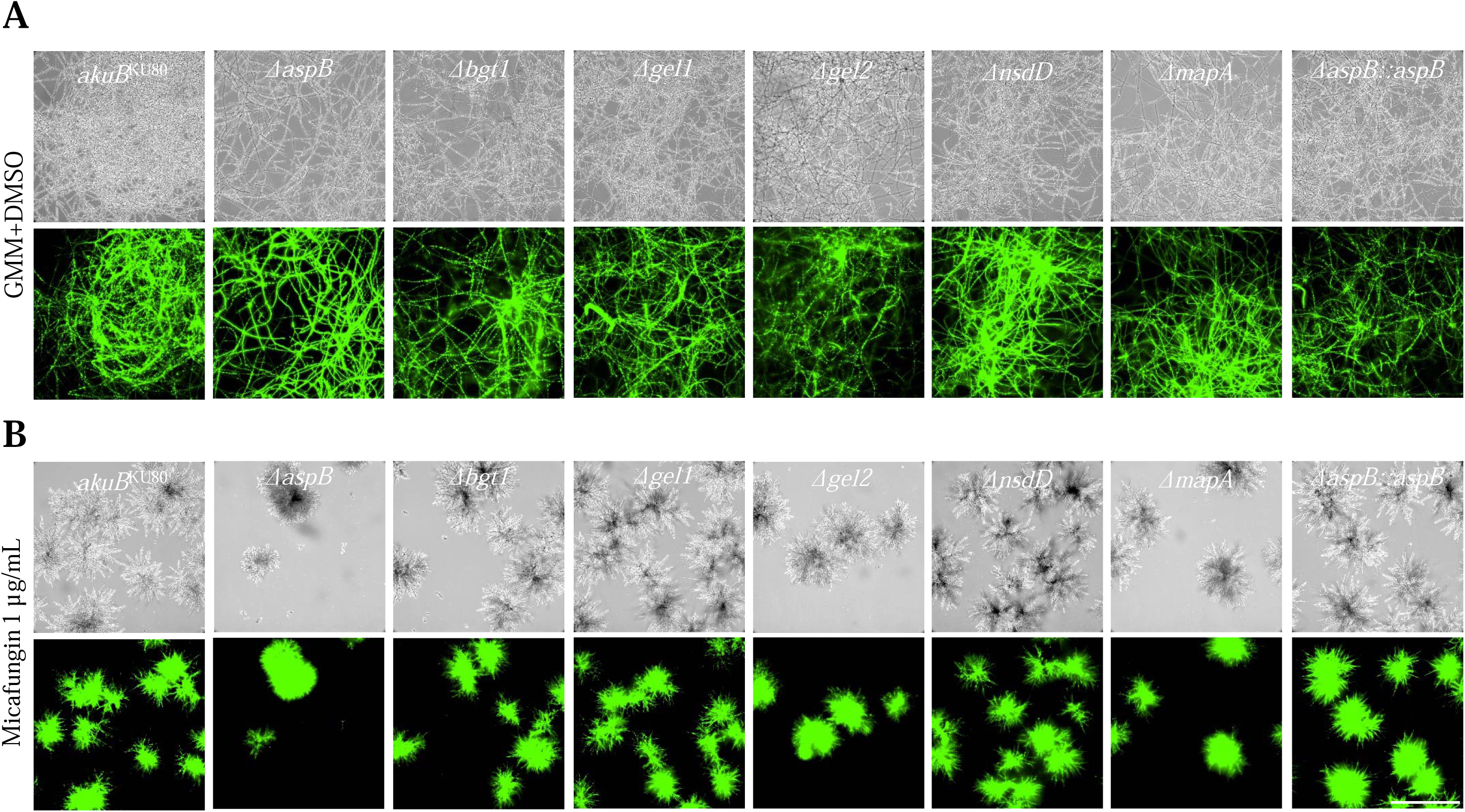
Candidate deletion mutants do not have reduced viability in micafungin. (A) All strains were fully viable when grown in GMM media supplemented with DMSO. Conidia (10^4^) were grown in GMM supplemented with 1 μL/mL DMSO for 24 hours on coverslips, then incubated in CFDA. Slides were prepared and visualized. (B) All strains have some viability when grown in micafungin. Conidia (10^4^) were grown on coverslips immersed in GMM supplemented with 1 μg/mL micafungin for 48 hours. Coverslips were then incubated in CFDA and visualized. Scale bar is 300 µm.

**Supplementary Figure 3.**
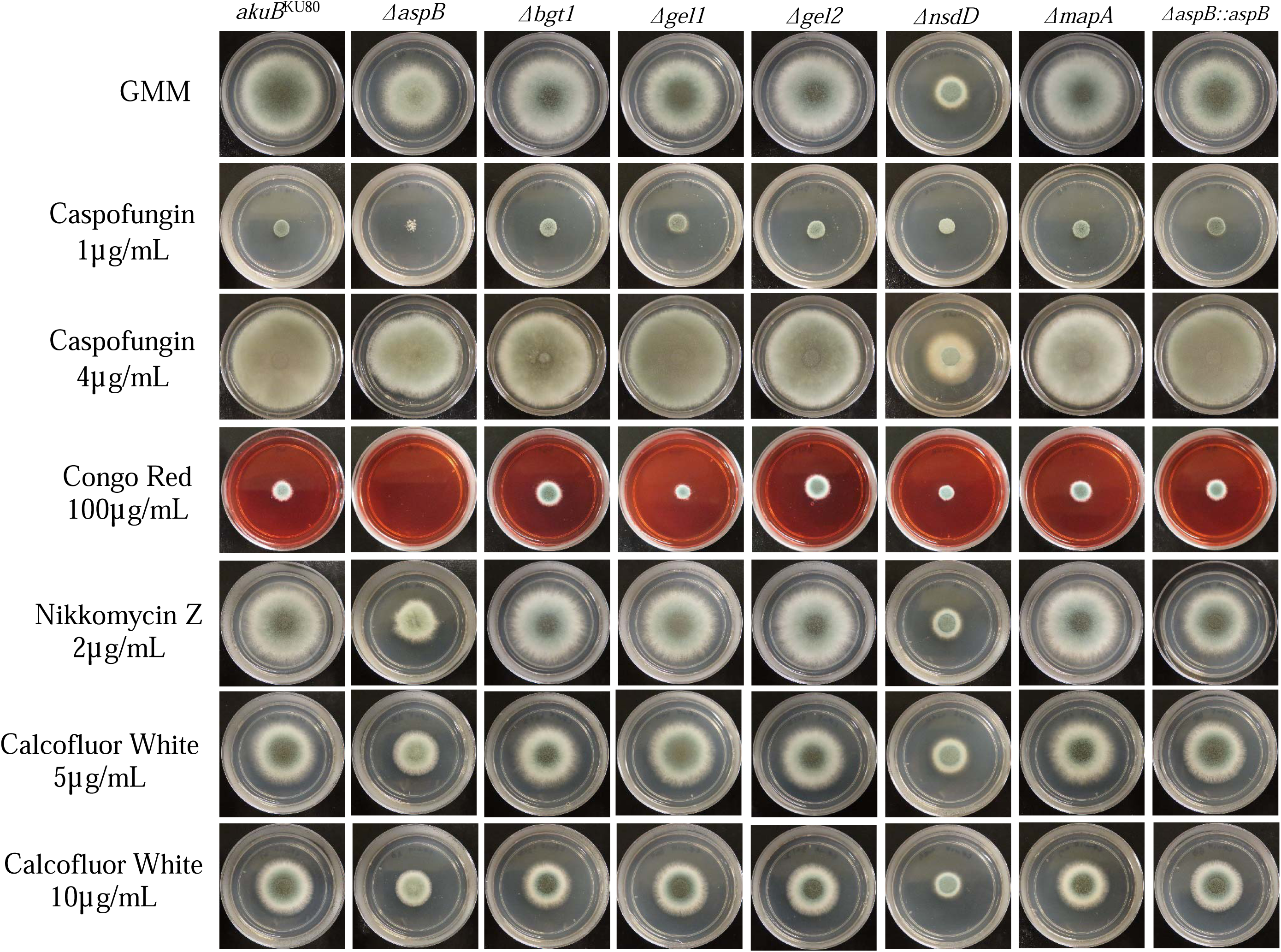
**Δ***bgt1*, **Δ***gel2*, and **Δ***nsdD* have mild sensitivity to calcofluorwhite. 10^4^ conidia were inoculated into GMM agar supplemented with cell wall disrupting agents listed in the figure then incubated for 3 days at 37°C, with the exception of 4 μg/mL caspofungin for five days to ensure observation of the caspofungin paradoxical effect. Experiments were replicated three times.

**Supplementary Figure 4.**
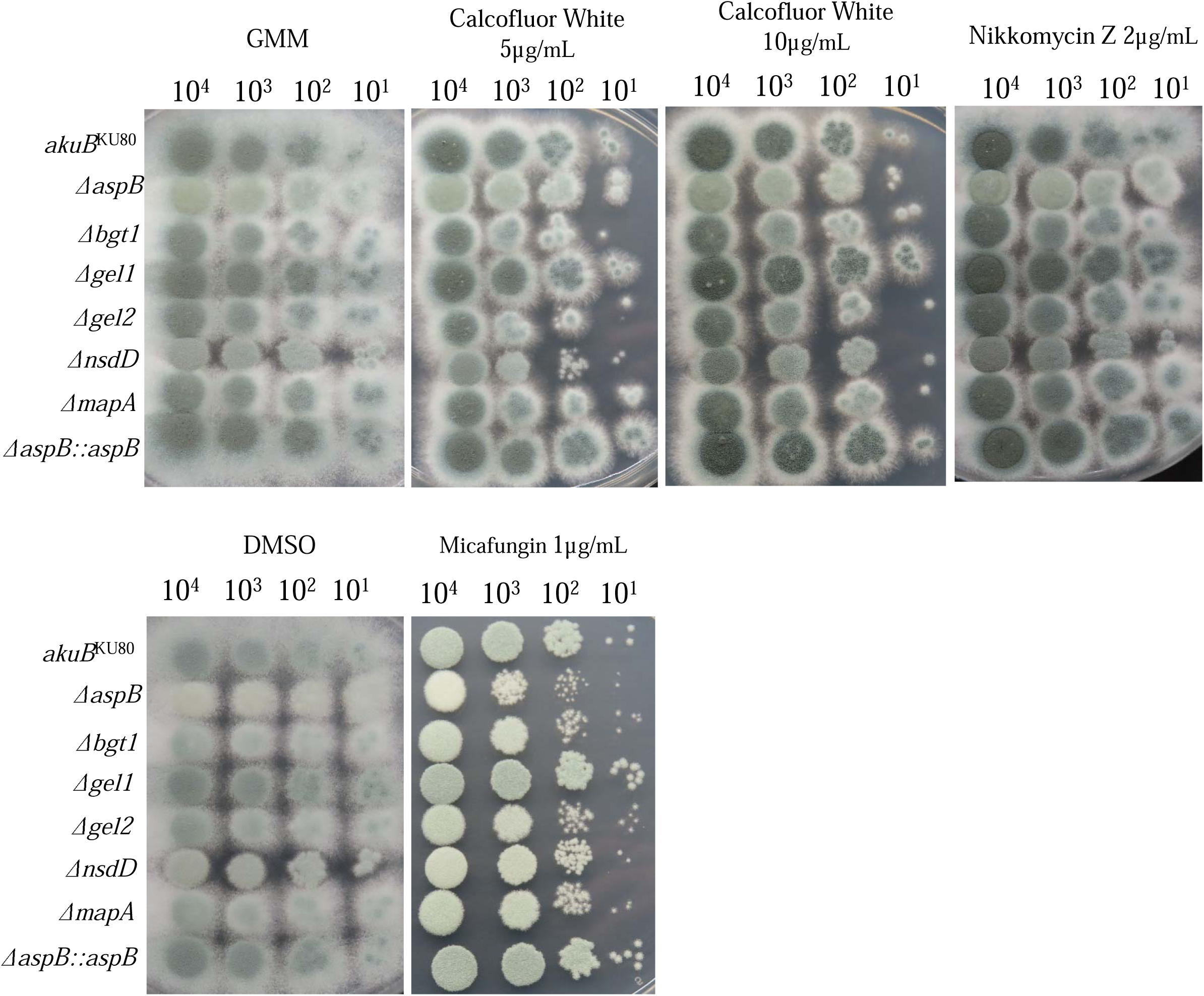
Candidate gene deletion mutants spot dilution assays confirm Δ*bgt1*, Δ*gel2*, and Δ*nsdD* sensitivity to calcofluor white. Conidia (10^4^-10^1^) were grown on GMM media supplemented with drug noted for 48 hours, then imaged. Experiments done in triplicate.

**Supplementary Figure 5.**
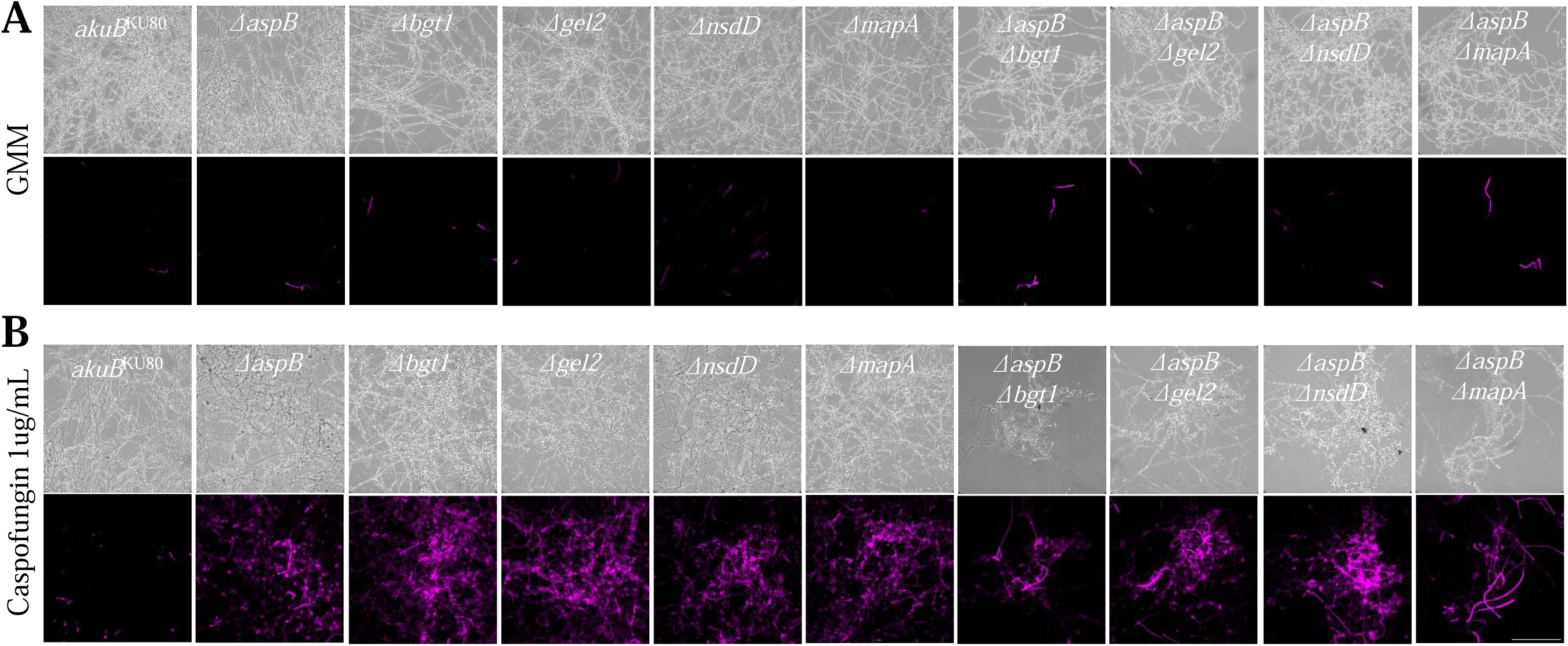
Double deletion mutants have hyphal damage after caspofungin exposure. (A) Double deletion mutants have minimal hyphal damage in basal conditions. Conidia (10^4^) were grown on coverslips in GMM for 24 hours. Coverslips were washed in PIPES (pH 6.7) for 5 minutes twice then treated with propidium iodide (PI) solution. Coverslips were washed two times with PIPES and visualized after. (B) Double deletion mutants have increased hyphal damage after caspofungin exposure. Conidia (10^4^) were grown on coverslips in GMM for 24 hours, then incubated in caspofungin for 2 hours. Cells were then washed with PIPES and incubated with PI. Scale bar is 250 µm. Experiments were replicated three times. All images were standardized in ImageJ.

